# BioRAG-DRAG: A Multimodal Biological Retrieval Layer for Local-First Biomedical Agents

**DOI:** 10.64898/2026.05.19.726174

**Authors:** Liang Wang

## Abstract

Biomedical agents need reliable access to heterogeneous evidence: literature text, gene and pathway records, protein sequences, DNA/cDNA sequences, and structured biological relations. Classical sequence tools such as BLAST remain the right choice for alignment-grounded verification, but they are not a unified context interface for large language model agents. We present **BioRAG-DRAG**, a local-first multimodal retrieval layer that combines pluggable neural sequence-text retrieval, BLAST verification, and graph-based evidence packaging. Specialized encoders such as ESM-2 can serve protein partitions, while OmniGene CPT provides a unified biological-language backbone for mixed sequence/text and agent-facing use; BLAST reranks or verifies sequence candidates; and DRAG graphs expose typed, traceable paths for downstream agents.

We introduce **BioRAG-Standard v0**, a partitioned corpus/library with 257,886 retrievable records and an initial annotation layer for engineering evaluation built from Open-Rosalind Standard biomedical records and sequence-window extensions. On an in-index sequence-window stress test, BLAST nearly saturates biological matching, while vector retrieval recovers substantial but lower biological match rates. On held-out parent-fragment controls, public protein encoders outperform the current OmniGene protein-window embedding, while DNA/cDNA dense retrieval remains weak even with off-the-shelf Nucleotide Transformer pooling; this supports a model-agnostic BioRAG design rather than a claim that one unified generator backbone is the best sequence-search encoder. Indexed Chroma lookup over Standard text and 100k sequence-window collections adds only small lookup overhead after query embedding; this does not measure end-to-end instant latency. Finally, exploratory sequence DRAG traces show inspectable biological neighborhoods, including immunoglobulin-family and gene-symbol modules, with initial graph controls indicating non-random but partly sequence-similarity-driven structure. These results support a bounded architecture: vector retrieval supplies unified candidate context, while BLAST and DRAG provide biological verification and evidence attribution.

## 1 Introduction

Biomedical information retrieval is inherently multimodal. A single user question may involve a gene symbol, a DNA fragment, a protein sequence, a pathway description, a variant, a paper abstract, or a combination of these. Traditional bioinformatics systems expose these modalities through separate tools: BLAST for local alignment [1], MMseqs2 or DIAMOND for high-throughput sequence search [2, 3], SQL or keyword search for database records, ontology lookup for GO terms, pathway databases for molecular processes, and graph traversal for relationships among entities. This division is scientifically useful, but it creates a practical gap for large language model (LLM) agents. Agents need a unified evidence interface that can retrieve, cite, and reason over heterogeneous local data without forcing the user or the model to manually choose a tool for each modality.

Retrieval-augmented generation (RAG) offers a natural interface for evidence grounding [4], but ordinary text RAG is not sufficient for biological data. DNA and protein sequences are not just strings of natural language, and exact sequence similarity has established statistical and biological foundations. Therefore, a biological RAG system should not frame neural vector retrieval as a replacement for BLAST. Instead, vector retrieval should act as a fast, shared candidate layer, while classical tools provide verification where their assumptions fit the query.

We propose **BioRAG-DRAG**, a multimodal biological retrieval layer for local-first biomedical agents. The core design is:

> query / sequence / mixed input
>
> →partitioned vector retrieval for coarse candidates and rapid context
>
> → BLAST verification or reranking for sequence-level evidence
>
> → DRAG graph expansion for typed evidence paths
>
> → local-first agent answer with citations and retrieval traces

This design separates three roles. First, vector retrieval provides a shared interface over text, DNA/cDNA, protein, and mixed records, while allowing different partitions to use different embedding models. Second, BLAST remains the biological alignment route for verification and reranking. Third, DRAG turns retrieval results into graph evidence that an agent can inspect, cite, and expand. The system is therefore complementary to BLAST rather than competitive with it.

The current empirical result supports this division as an engineering validation rather than a held-out homology benchmark. On the BioRAG-Standard v0 sequence-window stress test, vector retrieval is not as strong as full-database BLAST for alignment-grounded sequence verification, but it often places a biologically matched parent within a 200-candidate pool for downstream alignment-based verification. Held-out parent-fragment controls further show that protein-specialized ESM-2 and ProtT5 embeddings are stronger than the current OmniGene protein-window embedding, while the DNA/cDNA partition remains difficult for off-the-shelf dense pooling. These results reinforce the system claim: BioRAG should route partitions to appropriate encoders while preserving a unified agent evidence interface.

This paper makes four contributions:

1. **A reproducible local corpus/library export**. We build BioRAG-Standard v0 from Open-Rosalind Standard data and sequence-window extensions, yielding 257,886 corpus records and an initial annotation layer for engineering evaluation.
2. **A local-first hybrid retrieval architecture**. We implement vector coarse retrieval, BLAST verification, and DRAG graph packaging over Standard biomedical text, protein windows, DNA/cDNA windows, and mixed records.
3. **A retrieval and engineering evaluation**. We compare BLAST, vector retrieval, vector reranking, combined vector+BLAST evidence routes, held-out protein embedding baselines with ESM-2 and ProtT5, and an initial public DNA encoder control; we also decompose indexed lookup and verified-route latency.
4. **An exploratory DRAG evidence analysis**. We visualize typed sequence evidence traces and report biological-structure signals as hypothesis-generating results, with parent-collapse, k-mer graph, and degree-preserving null controls used to bound the interpretation.

## 2 Related Work

### Classical sequence search

BLAST remains a central tool for sequence similarity because it provides alignment-based scoring and interpretable statistical evidence [1]. Later systems such as MMseqs2 and DIAMOND improve speed and sensitivity at large scale, especially for protein and metagenomic workloads [2, 3]. This line of work is not a weak baseline for BioRAG; it is the biological verification layer that BioRAG should preserve. Our question is different: whether a neural multimodal retriever can provide a fast, shared candidate and context layer that cooperates with alignment search rather than replacing it.

### Biomedical language models and biomedical RAG

Domain-specific biomedical language models such as BioBERT, PubMedBERT, BioGPT, and BioMedLM improve biomedical text representation or generation by pretraining on biomedical corpora [5, 6, 7, 8]. RAG systems then combine model generation with retrieved evidence to reduce reliance on parametric memory [4]. Clinical and biomedical RAG systems such as Almanac and recent medical GraphRAG variants show the value of retrieval and graph evidence for safer medical question answering [9, 10]. BioRAG-DRAG is complementary to these systems: it focuses on local-first biological retrieval across text, DNA/cDNA, protein windows, and graph evidence, rather than on clinical QA alone.

### Biological foundation models and learned sequence search

Protein language models show that sequence-only pretraining can learn useful structure and function representations [11, 12, 13]. Dedicated protein-retrieval work such as PLMSearch demonstrates that learned protein representations can support fast remote-homology search [14]. More recently, ERAST combined large language models and vector database technology for scalable homology detection over approximately one billion biological sequences, covering both nucleotide and protein sequences [15]. Genomic language models, including DNABERT, DNABERT-2, Nucleotide Transformer, HyenaDNA, and Evo, similarly show that DNA-scale pretraining can produce transferable sequence representations [16, 17, 18, 19, 20]. These systems make clear that dense biological sequence search is not new by itself. BioRAG-DRAG instead studies how dense sequence-text retrieval can be packaged as a local biomedical-agent evidence layer together with BLAST verification, text records, and graph traces. We treat the embedding layer as pluggable: protein-only partitions can use public encoders such as ESM-2 or ProtT5, while OmniGene-4 CPT is useful as a unified biological-language backbone with expanded biological vocabulary for DNA, protein, structure-token, and natural-language inputs. Answer generation remains a separate agent/model function.

### Vector databases and graph-augmented retrieval

Dense vector retrieval has become a common substrate for RAG because approximate nearest-neighbor indexes can serve large precomputed embedding collections efficiently; FAISS is a representative GPU-capable system for billion-scale similarity search [21]. Graph-augmented RAG extends this substrate by organizing retrieved records and relations as paths or communities rather than isolated chunks; GraphRAG is one prominent recent example for text corpora [22]. Biological data are naturally graph structured: retrieved sequences often need to connect to genes, proteins, annotations, pathways, variants, and literature. BioRAG-DRAG evaluates both method-agnostic vector-neighbor graphs and BLAST-enriched hybrid graphs, asking whether biological structure appears before explicit domain rules are added.

### Local biomedical agents and this work’s lineage

BioRAG-DRAG builds on two local prerequisites. Open-Rosalind defines the auditable biomedical-agent contract: tool-mediated evidence, trace completeness, and bounded workflows [23]. OmniGene-4 provides a biological sequence-text foundation model: a Gemma-derived CPT model with expanded biological vocabulary for DNA, protein, structure tokens, and natural-language inputs [24]. This paper connects those directions by turning biological representations into a retrieval substrate and by packaging retrieval results into the evidence traces expected by a local biomedical agent; the retrieval layer itself remains model-agnostic.

## 3 BioRAG-Standard v0 Corpus

### 3.1 Corpus Construction

BioRAG-Standard v0 is built from the existing Open-Rosalind Standard index and local sequence-vector extensions. The exported dataset is stored under `data/biorag_standard_v0` and uses JSONL files for both corpus records and annotated tasks.

Each corpus row contains a stable dataset-local ID, source record ID, biological entity ID, source accession, modality, partition, retrievable text or sequence, extracted biological labels, and traceable metadata. Sequence-window records preserve parent accession, parent record ID, window offset, window size, stride, alphabet, and source FASTA header.

### 3.2 Annotated Tasks

The current annotation layer contains 112 retrieval tasks:

All tasks have at least one `positive_corpus_refs` link in the full export, enabling evaluation by direct corpus references as well as by expected labels such as entity IDs, source IDs, accessions, symbols, and biological gene labels.

### 3.3 Role in the Study

The corpus/library is the shared substrate for all retrieval conditions:

- text lookup over Standard records,
- vector search over Standard text and sequence windows,
- BLAST over sequence records,
- combined BioRAG retrieval with BLAST, vector, text, and graph routes,
- DRAG graph construction and biological enrichment analysis.

This shared substrate makes ablations cleaner: removing DNA, protein, text, graph, or BLAST routes changes the retrieval view without changing the underlying evidence universe. We do not claim that this v0 task layer is a comprehensive biological retrieval benchmark; it is an exported local evidence library with enough annotations to validate the first BioRAG workflow.

## 4 Method

### 4.1 Typed Multimodal Records

Records are typed before embedding or indexing. Examples include Standard biomedical text, protein sequence windows, DNA/cDNA sequence windows, and mixed English/sequence records. For sequence-window retrieval, the document body is the biological sequence itself, while metadata carries the parent accession and source information. This avoids embedding long header-heavy FASTA records as single documents and improves fragment retrieval.

The embedding layer is partitioned and pluggable. The main BioRAG-Standard sequence-window stress test uses the full merged OmniGene CPT model in BF16 so that DNA, protein, mixed sequence/text, and agent-facing biological-language inputs share one representation backbone. For protein-only held-out controls, we also build public ESM-2 and ProtT5 sequence-window indexes; for DNA/cDNA, we add an off-the-shelf Nucleotide Transformer 500M control. Embeddings are computed from the final hidden layer using attention-mask mean pooling over token hidden states followed by L2 normalization, not from next-token prediction logits. We also run last-token pooling or CLS-token pooling where supported as ablations. Windows use a size of 128 and stride of 64. The Chroma collections contain 100,000 DNA/cDNA windows and 100,000 protein windows for the main stress test; the controlled20k held-out protein control expands to 112,808 protein windows, and the controlled100k ProtT5 protein control expands to 589,210 windows.

### 4.2 Vector Coarse Retrieval

Vector retrieval serves two purposes:

1. **Instant mode**. It returns fast provisional context for interactive use.
2. **Candidate generation**. It produces top-*N* sequence/entity candidates for downstream biological verification.

For sequence-window queries, long inputs are split into query windows, then results are merged across windows. A lightweight sequence-aware reranker can optionally combine vector similarity with sequence overlap features. This reranker is not intended to replace BLAST; it is used to test whether simple local features can improve the vector candidate layer.

### 4.3 BLAST Verification and Reranking

For sequence inputs, BLASTP or BLASTN is used as the alignment-grounded verification route. In the current implementation, BLAST is run against local Swiss-Prot and Ensembl cDNA databases. The intended pipeline is:

~~~
vector top-N candidates
 -> BLAST fine scoring or full-database fallback
 -> reranked evidence with alignment metadata
~~~

We evaluate both full-database BLAST and candidate-subset BLAST. In the candidate-subset condition, vector retrieval first selects top-*N* parent accessions, `blastdbcmd` extracts those candidates from the local BLAST database, and BLASTP/BLASTN reranks only that candidate FASTA. This condition measures whether vector retrieval can supply a useful candidate set for verified BioRAG, while full-database BLAST remains the reference verification route.

### 4.4 DRAG Evidence Graphs

DRAG graphs convert retrieved records and relations into graph evidence. We build three sequence-window graph views:

- **vector-only graph:** edges are nearest-neighbor vector relations;
- **BLAST-only graph:** edges are alignment-derived sequence neighbors;
- **hybrid graph:** vector and BLAST edges are merged while preserving edge types.

The vector-only graph deliberately follows a method-agnostic text-RAG recipe: records become nodes, nearest-neighbor retrieval creates edges, and biological labels are used only for downstream analysis. This lets us test whether biological structure emerges before explicit biological rules are added.

## 5 Experiments

### 5.1 Retrieval Conditions

We evaluate the current local implementation as a workflow validation and stress test:

- **BLAST:** local BLASTP/BLASTN sequence retrieval;
- **Vector:** sequence-window Chroma retrieval using the configured embedding model;
- **Vector + sequence rerank:** vector retrieval plus a lightweight heuristic sequence-overlap reranker;
- **Combined BLAST+vector evidence:** BLAST and vector routes are both executed for sequence queries, with FTS, graph, and vector routes available for text and mixed records.

Metrics include Hit@*k*, MRR, biological Hit@*k*, biological MRR, average latency, route coverage, and retrieval traces.

The 100-query sequence-window set is not a held-out homology benchmark: queries are fragments or perturbations derived from indexed parent sequences. We therefore use it to test local retrieval behavior, candidate-pool quality, and evidence packaging. Separately, we evaluate a 500-query held-out parent-fragment control over a controlled20k protein index to compare BLAST, OmniGene, and public ESM-2 embeddings under a more realistic no-exact-parent setting.

### 5.2 Engineering Latency

Latency is decomposed into vector-index lookup, BLAST route time, graph expansion, verified-mode median sum, and FAISS CPU lookup. All vector lookup measurements exclude model cold start, query embedding generation, LLM generation, and network calls. This isolates the indexed retrieval stage after a resident embedding service has produced the query vector.

### 5.3 Graph Evidence Structure

We evaluate whether DRAG can package retrieved sequence records into typed, inspectable graph neighborhoods. The main text reports a qualitative evidence-trace visualization and a compact graph-control summary. Community purity, GO/Reactome enrichment, and PubMed-sharing analyses are treated as exploratory appendix results because they remain underpowered for biological mechanism claims.

## 6 Results

### 6.1 Retrieval Quality

On the 100-query BioRAG-Standard sequence-window stress test, the retrieval results are shown in Table 3. BLAST remains the strongest verification route. Vector-only retrieval is weaker on exact parent-ID recovery, but it recovers biologically related neighborhoods often enough to serve as a useful coarse-retrieval layer in this local setting. The heuristic sequence-aware reranker narrows much of the gap; we treat this as an engineering ablation rather than a methodological contribution. The combined BLAST+vector evidence route preserves BLAST-level biological recall while retaining vector neighborhoods and route traces for agent use. These results should not be interpreted as evidence of held-out biological retrieval competitiveness.

**Table 1:** BioRAG-Standard v0 corpus partitions.

| Corpus partition | Records | Source |
| --- | --- | --- |
| Standard text | 57,856 | HGNC genes, GO terms, Reactome pathways, ClinVar gene summaries |
| Protein sequence windows | 100,000 | Swiss-Prot windows embedded with OmniGene CPT |
| DNA/cDNA sequence windows | 100,000 | Ensembl cDNA windows embedded with OmniGene CPT |
| Mixed POC records | 30 | mixed English/sequence examples |
| **Total** | **257,886** | text + protein + DNA/cDNA + mixed |

**Table 2:** BioRAG-Standard v0 annotation families.

| Task family | Records |
| --- | --- |
| Gene lookup | 5 |
| Pathway lookup | 3 |
| Protein sequence retrieval | 52 |
| DNA/cDNA sequence retrieval | 52 |
| **Total** | **112** |

**Table 3:**
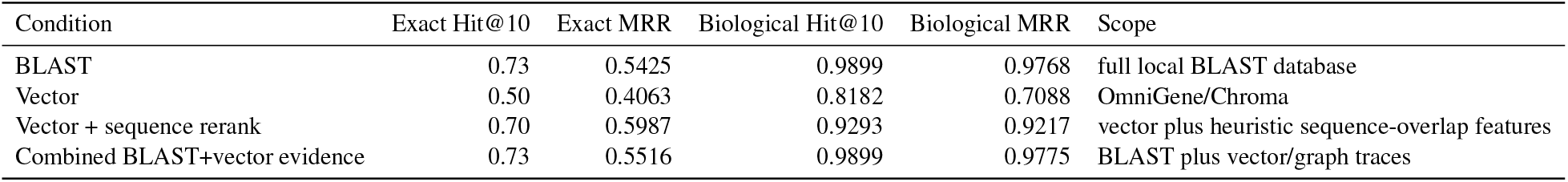
Retrieval quality on the 100-query BioRAG-Standard sequence-window stress test. This evaluates local fragment recovery and candidate-pool behavior, not held-out biological homology search.

As a candidate-pool ablation, vector-to-candidate-BLAST reranking reaches biological Hit@10/MRR of 0.9293/0.9293 with 200 vector-selected parent candidates, with candidate biological Recall@200 of 0.9596. This shows that, in this in-index local stress test, vector retrieval often places the biologically matched parent within a 200-candidate pool for downstream alignment-based verification. It does not establish a speed advantage; matched 100k/300k/full vector scale curves are required before candidate-BLAST can be claimed as a systems improvement. The full budget sweep is therefore reported only in Appendix A.

The interpretation is therefore not “vector beats BLAST.” The result supports a two-stage engineering strategy: vector retrieval supplies unified candidates and provisional context; BLAST supplies alignment-grounded verification.

### 6.2 Held-Out Protein Embedding Baselines

To separate BioRAG system design from the choice of embedding backbone, we also run a 500-query held-out parent-fragment control on a controlled20k protein index. The exact held-out parent accessions are excluded from the index; the main metric is whether retrieval finds an indexed biologically related record. Table 4 shows that BLAST remains the strongest alignment-grounded route, while public protein encoders are substantially stronger than the current OmniGene protein-window embedding on this protein-only retrieval task. ProtT5 mean pooling is the strongest completed dense baseline. Last-token pooling is weak across OmniGene, ESM-2, and ProtT5, so mean pooling remains the default embedding mode.

**Table 4:** Held-out parent-fragment protein retrieval on the controlled20k index. Vector rows use sequence-window Chroma retrieval with a top-200 candidate pool.

| Method | Pooling | Biological Hit@10 | Biological MRR | Recall@200 | Mean latency |
| --- | --- | --- | --- | --- | --- |
| BLAST | alignment | 0.8560 | 0.8289 | 0.8560 | 138.6 ms |
| OmniGene-4-CPT BF16 | mean | 0.3120 | 0.2623 | 0.4940 | 538.4 ms |
| OmniGene-4-CPT BF16 | last-token | 0.0940 | 0.0725 | 0.1840 | 531.5 ms |
| ESM-2 650M | mean | 0.6560 | 0.5764 | 0.7340 | 329.0 ms |
| ESM-2 650M | last-token | 0.2360 | 0.2025 | 0.3560 | 332.2 ms |
| ProtT5-XL-UniRef50 | mean | 0.7780 | 0.7158 | 0.8460 | 329.1 ms |
| ProtT5-XL-UniRef50 | last-token | 0.4680 | 0.3668 | 0.6400 | 333.1 ms |

**Table 5:** Held-out DNA/cDNA parent-fragment control. BLASTN is the alignment-grounded reference. Vector rows use top-200 Chroma retrieval; the Nucleotide Transformer row indexes the full controlled20k window set.

| Method | Index setting | Biological Hit@10 | Biological MRR | Mean latency |
| --- | --- | --- | --- | --- |
| BLASTN | controlled20k cDNA | 0.9100 | 0.9100 | 94.7 ms |
| OmniGene-4-CPT BF16 mean | controlled20k windows | 0.1000 | 0.0795 | 1049.0 ms |
| Nucleotide Transformer 500M mean | full controlled20k windows | 0.0100 | 0.0021 | 992.5 ms |
| BLASTN | full local cDNA | 0.9100 | 0.9100 | 328.3 ms |

This result changes the interpretation of OmniGene in BioRAG. OmniGene should not be presented as the strongest protein-only dense retriever in the current experiments, and last-token pooling is a poor choice for this fragment-retrieval setting. Its role is better framed as a unified biological-language backbone for mixed sequence/text and agent workflows. BioRAG can use specialized encoders such as ProtT5 or ESM-2 for protein partitions while keeping the same Chroma, BLAST, DRAG, and citation interfaces.

### 6.3 Held-Out DNA/cDNA Control

We also construct a 100-query DNA/cDNA parent-fragment control from Ensembl cDNA. Exact held-out transcript accessions are removed from the index, and biological matching uses shared gene symbol labels. To make the small vector-index experiment interpretable, we build a controlled20k cDNA subset containing at least one non-held-out same-gene transcript for every query plus random background records. This split has zero exact held-out transcript leakage, but 62 of 100 queries are exact substrings of other indexed transcripts, reflecting isoform and conserved-fragment overlap; we therefore treat it as a held-out transcript-fragment control rather than a strict remote-homology benchmark.

This DNA result is intentionally reported because it prevents overclaiming. The current OmniGene sequence-window embedding is weak on this DNA-only transcript-fragment retrieval control, whereas BLASTN remains strong. Nucleotide Transformer 500M does not improve the result under off-the-shelf mean pooling, and a smaller 20k-window CLS-pooling smoke run also remains low at 0.0300/0.0090 Bio Hit@10/MRR. We therefore do not conclude that dense DNA retrieval is solved by simply swapping in a genomic foundation model. The result reinforces the central system design: BioRAG should keep a unified evidence interface while allowing the DNA partition to use retrieval-specific genomic encoders, fine-tuning, or alignment verification.

### 6.4 Engineering Latency

Standard text and 100k BF16 sequence-window indexes give the lookup and verification times in Table 6. These results support a **lookup/verified** product design. After a query embedding is available from a resident embedding service, vector lookup adds only a small indexed-search overhead; in verified mode, BLAST and DRAG add biological grounding and evidence attribution. We do not report end-to-end instant latency here because query embedding time is excluded. The current FAISS GPU wheel detects the RTX PRO 6000 Blackwell GPU but lacks compatible sm 120 kernels; GPU lookup is therefore left as an engineering follow-up rather than a reported result.

**Table 6:** Indexed lookup and verified-route latency.

| Component | Text / Standard | DNA/cDNA | Protein | Notes |
| --- | --- | --- | --- | --- |
| Chroma lookup, top-10 | 4.6924 ms | 5.9630 ms | 6.0948 ms | query embedding excluded |
| BLAST top-10 | n/a | 294.5684 ms | 1081.9300 ms | local BLAST |
| Hybrid graph expansion | n/a | 0.3540 ms | 0.3254 ms | SQLite graph expansion |
| Verified vector+BLAST+graph | n/a | 300.8221 ms | 1088.8706 ms | median route-latency sum |
| FAISS CPU lookup, IndexFlatIP | 8.3346 ms | 15.4626 ms | 15.6090 ms | flat top-10 lookup |

### 6.5 Swiss-Prot Scale Frontier

We further measure the scale frontier on the held-out protein parent-fragment benchmark. Starting from the full held-out Swiss-Prot-derived index, we create controlled 100k and 300k subsets that preserve same-gene positive candidates for all 500 queries and have zero exact held-out-parent leakage. Table 7 reports completed BLAST references, completed ProtT5 mean vector points at controlled20k, controlled100k, and controlled300k, and vector-to-candidate-BLAST reranking points at controlled100k and controlled300k. The full-scale dense index remains pending.

**Table 7:** Swiss-Prot protein scale frontier on the 500-query held-out parent-fragment benchmark. Vector rows use ProtT5 mean pooling with top-200 Chroma retrieval. Candidate-BLAST is reported as an ablation, not as a speed advantage in the current Chroma implementation.

| Scale | Records | BLAST Bio@10 | BLAST MRR | BLAST ms | Vector Bio@10 | Vector MRR | Vector R@200 | Vector ms | Cand. Bio@10/MRR | Cand. ms |
| --- | --- | --- | --- | --- | --- | --- | --- | --- | --- | --- |
| controlled20k | 20,000 | 0.8560 | 0.8289 | 138.6 | 0.7780 | 0.7158 | 0.8460 | 329.1 | — | — |
| controlled100k | 100,000 | 0.8320 | 0.7960 | 329.2 | 0.7260 | 0.6268 | 0.8000 | 581.4 | 0.7580/0.7245 | 839.7 |
| controlled300k | 300,000 | 0.8140 | 0.7613 | 796.5 | 0.6760 | 0.5788 | 0.7740 | 812.5 | 0.7320/0.6835 | 1020.6 |
| full held-out Swiss-Prot | 483,581 | 0.8060 | 0.7442 | 1039.3 | — | — | — | — | pending | — |

This table is a boundary result rather than a speed claim. BLAST remains the verified reference. ProtT5 mean vector retrieval provides useful but lower dense retrieval, with biological Recall@200 of 0.8460, 0.8000,and 0.7740 at controlled20k, controlled100k, and controlled300k. Candidate-BLAST improves MRR over vector-only at controlled100k (0.7245 vs. 0.6268) and controlled300k (0.6835 vs. 0.5788), but measured total latency remains slower than full-database BLAST at both matched scales. The candidate-BLAST-only portions are much smaller, 189.2 ms and 209.6 ms after vector candidates are available, so the systems bottleneck in this Chroma POC is vector candidate serving rather than candidate alignment. Thus the current result supports candidate-BLAST as an evidence-quality and routing ablation; a systems speed advantage remains a larger-scale or optimized vector-serving hypothesis.

### 6.6 DRAG Evidence Graph Showcase

Figure 5 shows two real DRAG traces. The DNA/cDNA example centers on an IGKV sequence neighborhood, while the protein example centers on a yfbR neighborhood. In both cases, the graph view exposes which edges come from vector similarity and which come from BLAST-derived alignment evidence. This is the main agent-facing value of DRAG in the current paper: retrieved biological sequences are not only ranked, but packaged as typed neighborhoods that can be inspected, cited, and expanded.

**Figure 1:**
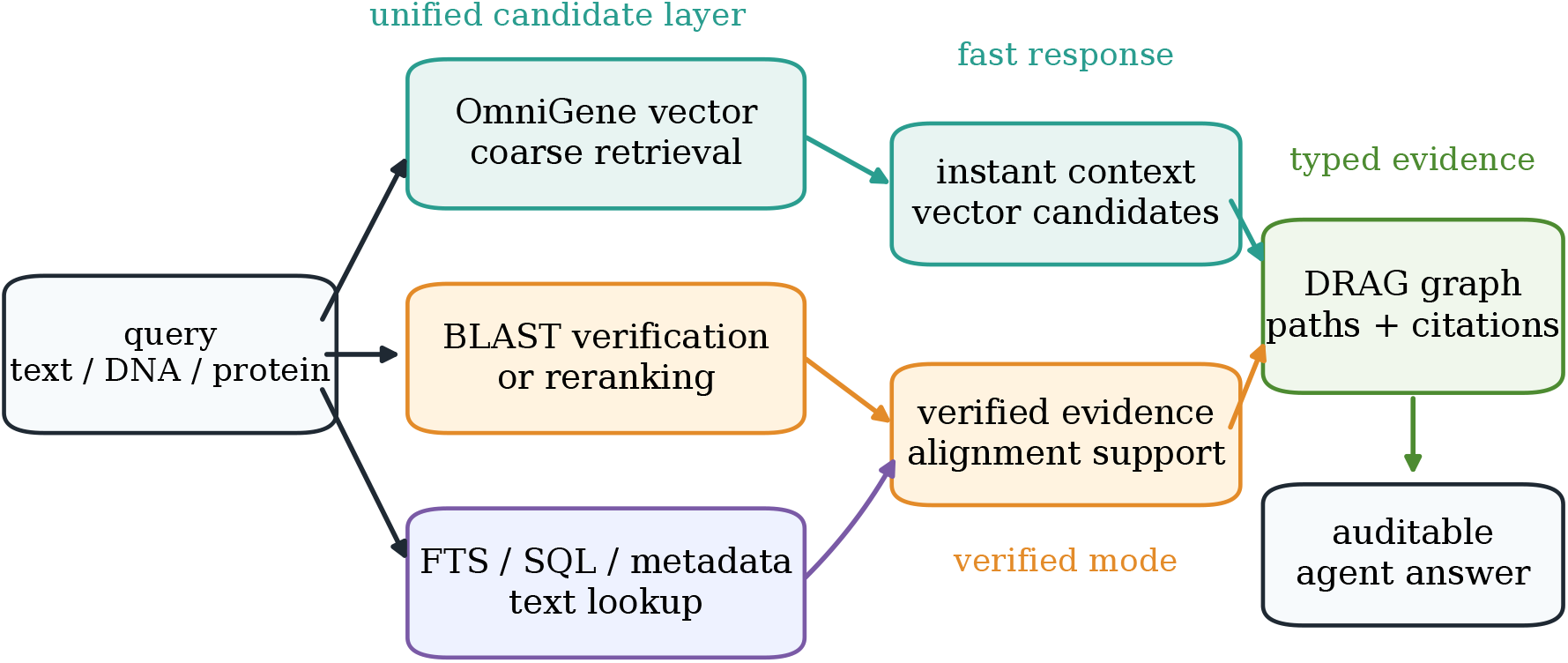
BioRAG-DRAG architecture. Vector retrieval provides a rapid multimodal candidate layer after query embedding; BLAST supplies sequence verification and reranking; DRAG packages vector, alignment, text, and graph evidence into typed traces for a local biomedical agent.

**Figure 2:**
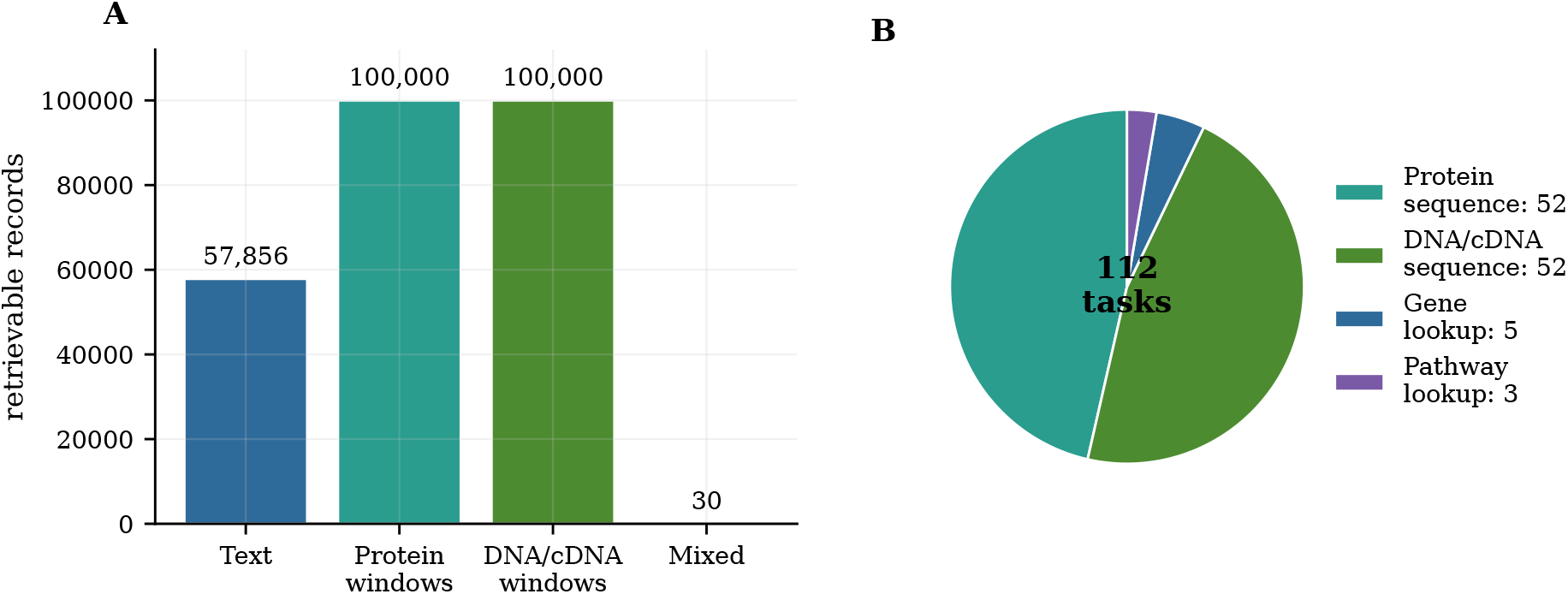
BioRAG-Standard v0 composition. The corpus/library contains 257,886 retrievable records across Standard text, protein windows, DNA/cDNA windows, and mixed records, with a first 112-task annotation layer.

**Figure 3:**
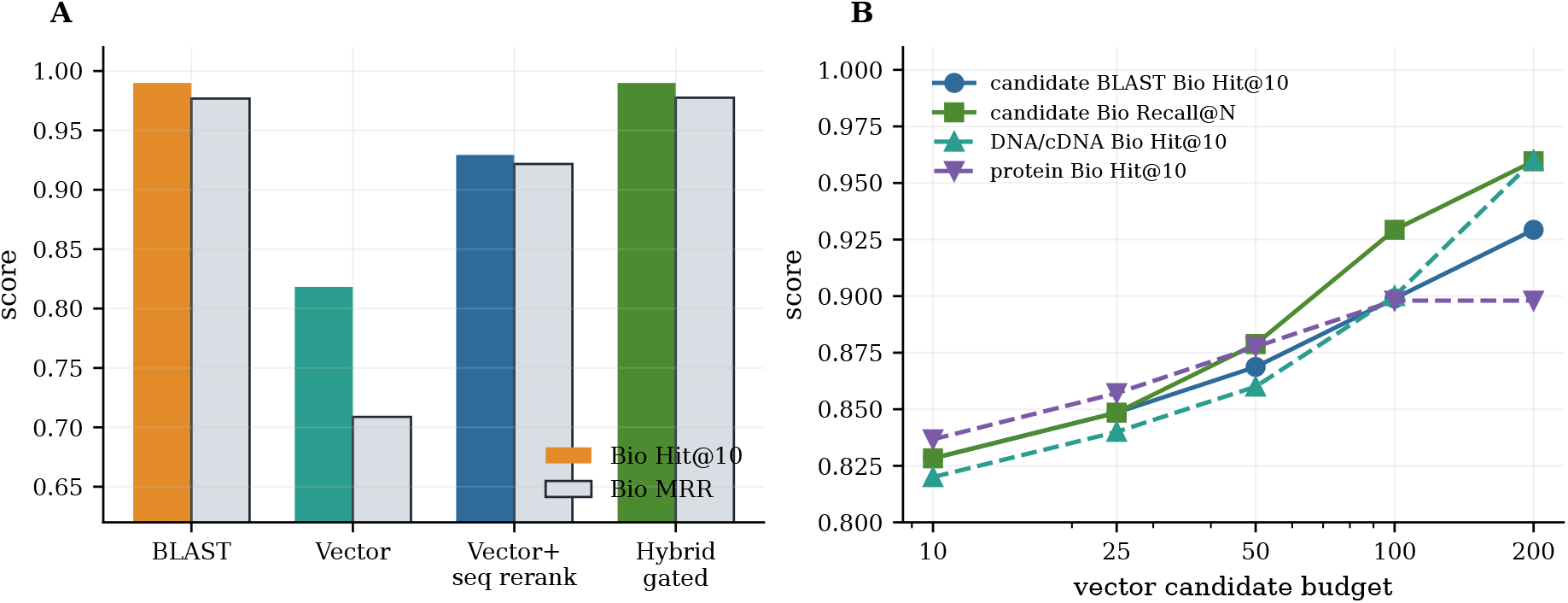
Retrieval quality on the 100-query sequence-window stress test. BLAST remains the strongest alignment-grounded verifier, while vector retrieval supplies lower but useful local candidate context; candidate-BLAST is treated as an ablation rather than a peer condition.

**Figure 4:**
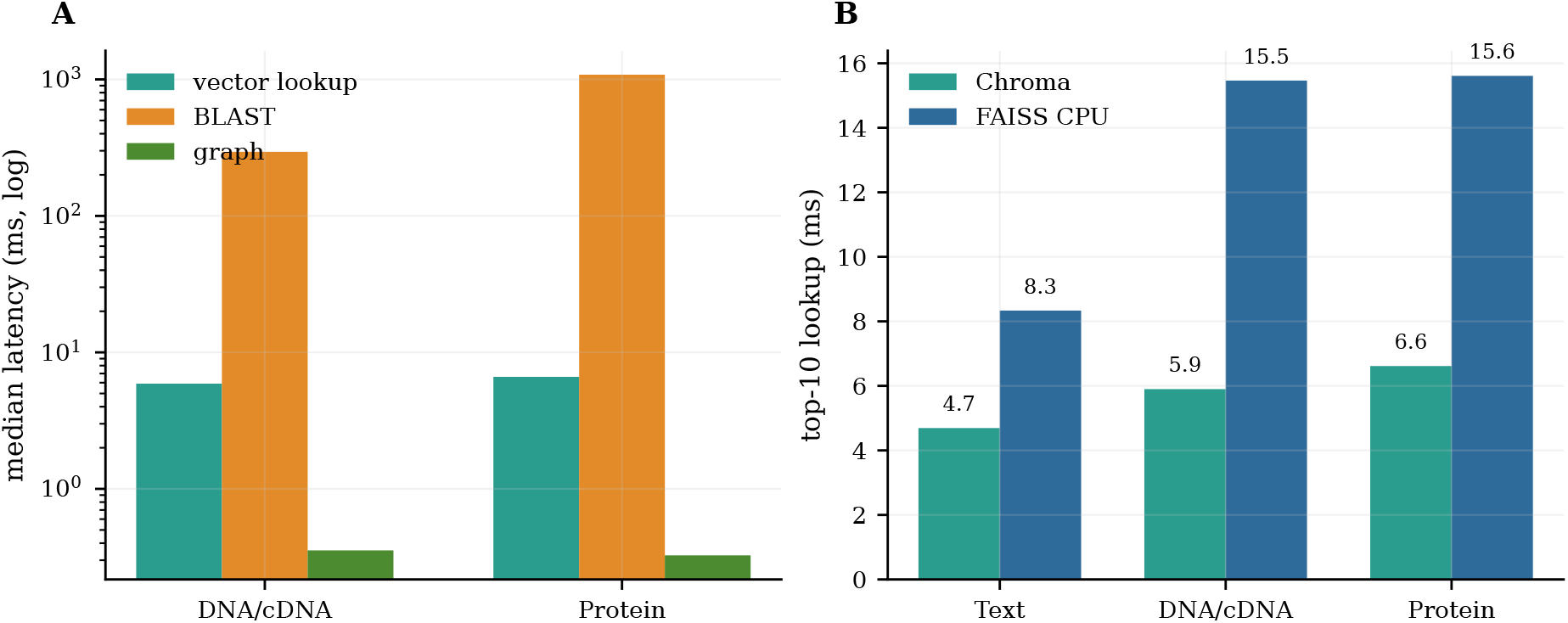
Indexed lookup and verified-route latency. Vector lookup is measured after query embedding; BLAST dominates verified-mode latency, while graph expansion is negligible.

**Figure 5:**
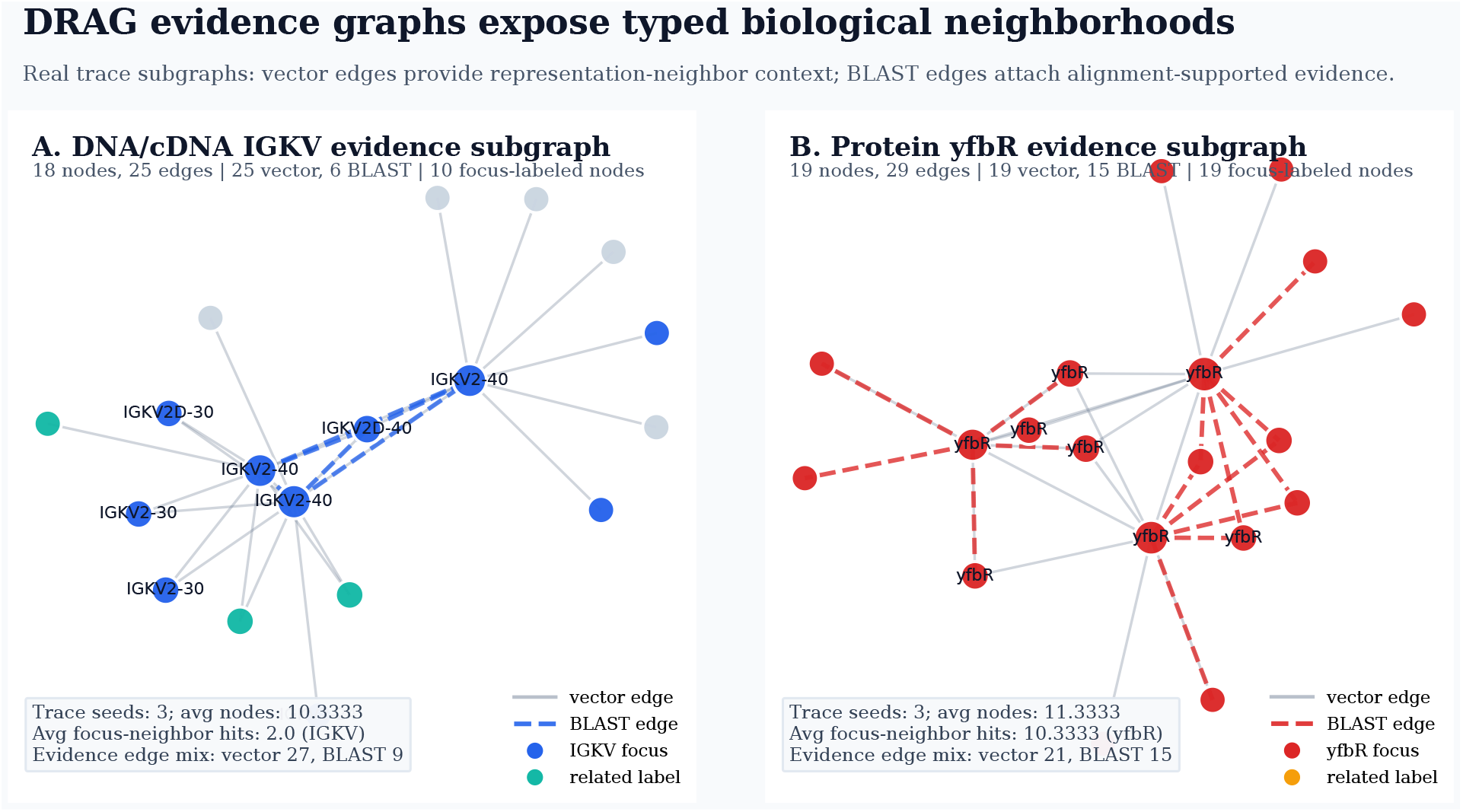
DRAG evidence graph showcase from real local traces. Panel A shows an IGKV-focused DNA/cDNA subgraph; panel B shows a `yfbR`-focused protein subgraph. Solid gray edges are vector-neighbor evidence and dashed colored edges are BLAST-neighbor evidence. The figure illustrates the agent-facing value of DRAG: retrieved biological sequences are packaged as typed, inspectable neighborhoods rather than isolated ranked hits.

**Figure 6:**
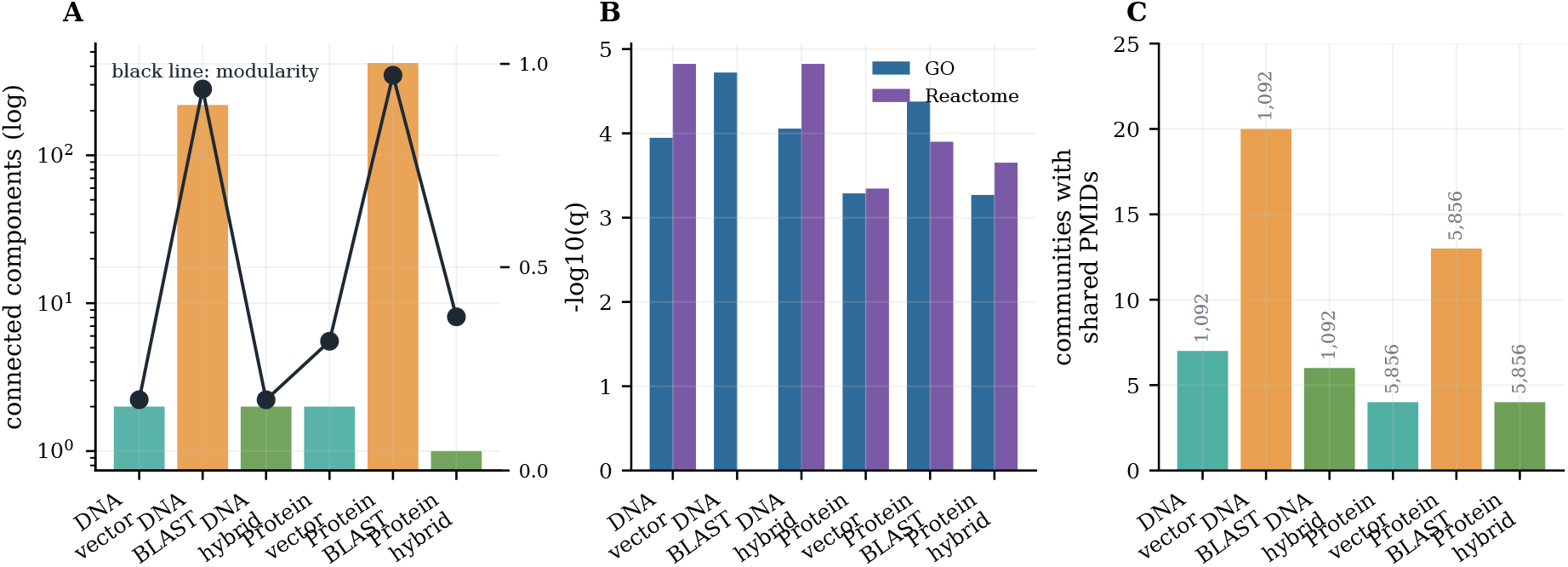
Exploratory DRAG biological-structure analysis. Vector, BLAST, and hybrid graphs expose different graph connectivity, functional enrichment, and literature-sharing patterns. These signals support hypothesis generation and should be interpreted with the parent-collapse, k-mer/Jaccard, and degree-preserving null controls in Table 8.

These examples are exploratory. Appendix B reports community purity, GO/Reactome enrichment, and shared-PubMed analyses. The controls in Table 8 support non-random graph structure, but they also show that simple sequence similarity explains part of the signal.

**Table 8:** Compact DRAG graph controls on 10k graph views. Existing DRAG views are already parent-collapsed. Observed graphs show label-enriched communities and degree-preserving null graphs do not, but k-mer/Jaccard graphs also recover strong sequence-family signals; therefore the biological interpretation remains exploratory.

| Modality | Condition | Nodes | Edges | Communities | Top label signal |
| --- | --- | --- | --- | --- | --- |
| DNA/cDNA | observed vector graph | 579 | 19,344 | 9 | biotype:IG_V_gene, 21 nodes, purity 1.00 |
| DNA/cDNA | k-mer/Jaccard graph | 579 | 2,111 | 26 | gene family:IGKV, 29 nodes, purity 1.00 |
| DNA/cDNA | degree-preserving null | 579 | 19,344 | 4.2 | no comparable top signal |
| Protein | observed vector graph | 2,623 | 31,794 | 16 | gene symbol:nbaC, 7 nodes, purity 0.50 |
| Protein | k-mer/Jaccard graph | 2,623 | 9,398 | 107 | gene symbol:pgl, 28 nodes, purity 1.00 |
| Protein | degree-preserving null | 2,623 | 31,794 | 12.6 | no comparable top signal |

### 6.7 Agent Evidence Case Study

To test the agent-facing evidence contract without adding another generator model, we convert existing DRAG traces into answer-ready evidence packs. No LLM is called in this experiment. The case-study script summarizes the graph context that a downstream agent would receive, including route labels, representative edges, and an explicit boundary on what may be claimed from the available evidence.

Table 9 shows the operational distinction between retrieval routes. For the IGKV2-40 case, the hybrid pack contains BLAST-supported cDNA neighbors with 100% identity as well as high-score vector neighbors from the same immunoglobulin family. For the protein pgl and yfbR cases, vector traces recover broad same-label neighborhoods, while BLAST edges provide alignment-labeled support for citation. The resulting answer skeleton is deliberately conservative: the agent may describe the retrieved neighborhood and cite whether each edge is vector-derived or BLAST-derived, but it should not infer function, mechanism, clinical relevance, or experimental conclusions unless annotation, pathway, domain, or literature evidence is also retrieved.

**Table 9:**
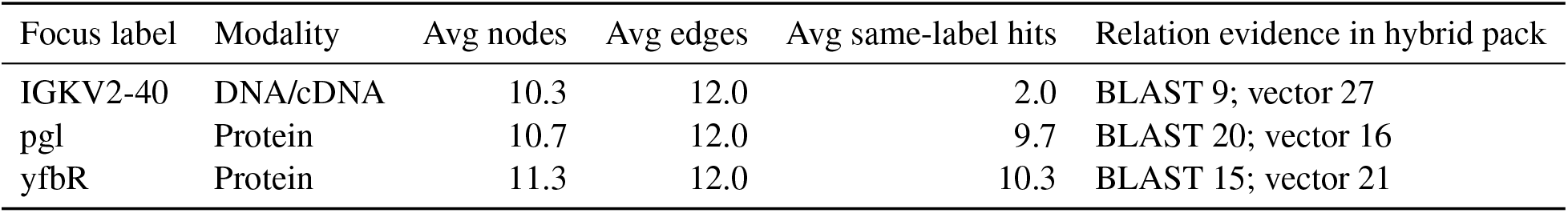
Agent-style evidence packs generated from DRAG traces. Hybrid DRAG keeps both broad vector context and BLAST-labeled verification edges visible to the downstream agent.

| Focus label | Modality | Avg nodes | Avg edges | Avg same-label hits | Relation evidence in hybrid pack |
| --- | --- | --- | --- | --- | --- |
| IGKV2-40 | DNA/cDNA | 10.3 | 12.0 | 2.0 | BLAST 9; vector 27 |
| pgl | Protein | 10.7 | 12.0 | 9.7 | BLAST 20; vector 16 |
| yfbR | Protein | 11.3 | 12.0 | 10.3 | BLAST 15; vector 21 |

## 7 Discussion

### 7.1 Why Vector Retrieval Matters Even When BLAST Is Stronger

BLAST is stronger for alignment-grounded verification and should remain the baseline for exact sequence similarity. However, agent systems need more than exact sequence matching. They need a unified evidence interface that can retrieve text, sequences, annotations, and graph paths, then package them into prompt-ready citations. Vector retrieval provides this shared substrate. The held-out ESM-2 and ProtT5 results further suggest that BioRAG should not force all partitions through one embedding model: specialized protein encoders can be used where they are stronger, while OmniGene remains useful for unified biological-language and mixed-modality contexts. The right comparison is not vector versus BLAST as mutually exclusive tools, but vector as coarse candidate generation and BLAST as fine verification.

### 7.2 Lookup and Verified Retrieval Modes

The latency results motivate two operating modes. Lookup mode returns vector-only context after query embedding and is useful for rapid local evidence preview. Verified mode runs BLAST and DRAG expansion to attach biologically grounded evidence. This design fits biomedical agents: the user can receive provisional context from a unified local index, while the system appends verified evidence when the slower routes finish. A full end-to-end latency claim requires warm embedding measurements and is left for the next experimental pass.

### 7.3 DRAG as a Biological Representation Probe

The graph results suggest that sequence-vector neighborhoods may contain biological structure, not just retrieval convenience. The current main-text claim is intentionally narrow: DRAG provides an inspectable evidence package for agents, and its sequence communities are promising exploratory objects. Initial controls sharpen this boundary. The analyzed graph views are already parent-collapsed, and degree-preserving null graphs do not show comparable label-purity signals, which argues against pure topology as the explanation. However, k-mer/Jaccard graphs also recover strong family-level communities, so part of the observed DRAG signal is explained by ordinary sequence similarity. Stronger biological-discovery claims require larger parent-collapsed graphs and domain, GO/pathway, and literature case studies against these controls.

## 8 Limitations

BioRAG-Standard v0 is still a first version. The current tasks emphasize lookup and sequence-fragment retrieval, and the main 100-query sequence test uses fragments or perturbations from indexed parent sequences. It should therefore be interpreted as a local workflow stress test rather than a held-out biological homology benchmark. Multi-hop sequence-to-function, sequence-to-pathway, and literature-grounded tasks are needed for stronger agent evaluation. The mixed partition is small and should be expanded with abstracts, UniProt function text, pathway descriptions, and sequence snippets.

The current vector reranker is lightweight and heuristic, and candidate-subset BLAST reranking is limited by whether the vector candidate pool contains the correct or biologically equivalent parent sequence. A preliminary graph-expanded retrieval ablation used 10k view graphs rather than a full 100k or complete sequence graph; it added graph candidates but did not improve recall beyond the vector seed pool, so it is treated as a coverage-limited result rather than a central claim. FAISS GPU lookup is not reported because the available wheel lacks Blackwell-compatible kernels on the current RTX PRO 6000 machine. Functional and literature enrichment coverage depends on local UniProt/NCBI/HGNC mappings; Pfam or InterPro domain enrichment and title-level literature case studies would strengthen biological interpretation.

The embedding comparison is still incomplete. We added ESM-2 and ProtT5 protein baselines plus a mean-vs-last pooling ablation, and we added an initial DNA/cDNA held-out transcript-fragment control with Nucleotide Transformer 500M as a public DNA encoder control. However, newer ESM-family checkpoints and original Gemma4 MoE controls remain to be evaluated, and DNABERT-2 is currently blocked by custom-model compatibility issues under the installed Transformers/Torch environment; it should be rerun in a pinned Transformers 4.x environment before submission if DNA public baselines become central. The current DNA control suggests that neither OmniGene nor off-the-shelf Nucleotide Transformer pooling should be treated as a strong DNA-only retriever in this setting. This separates the contribution of the unified BioRAG system from the contribution of biological tokenization, continued pretraining, and partition-specialized encoders. The BLAST side of the Swiss-Prot scale curve and the controlled100k/300k ProtT5 vector/candidate-BLAST points are now measured, but full-scale vector indexes, FAISS/Milvus-style optimized serving, and candidate-subset BLAST sweeps are still needed before claiming candidate-BLAST speed advantages.

The biological-meaning analysis should also be interpreted carefully. Community purity, GO/Reactome enrichment, and shared PubMed IDs show that DRAG neighborhoods are non-random and inspectable, but they do not prove causal mechanisms or validate novel wet-lab hypotheses. The appropriate use is hypothesis generation, evidence organization, and agent grounding. Biological claims that affect experimental design, diagnosis, treatment, or safety-critical decisions should still be verified by domain experts and specialized tools.

## 9 Reproducibility and Use Scope

The implementation is local-first and will be released at https://github.com/maris205/biorag. The corpus export lives under `data/biorag_standard_v0;` retrieval and graph reports are stored under reports/; and the main evaluation scripts are in `scripts`. The main BioRAG sequence-vector experiments use the full OmniGene-4 CPT merged model in BF16, not the GGUF or 4-bit quantized variants, while the held-out protein baselines use ESM-2 and ProtT5 BF16 embeddings. The vector store is Chroma for the POC because it supports simple CRUD and collection management, while FAISS CPU is used as a lightweight lookup-speed reference. BLAST runs against local Swiss-Prot and Ensembl cDNA databases.

The main reported commands are documented in `docs/REPRODUCIBILITY.md,` the evaluation notes, and the research matrix, including vector candidate-budget sweeps, vector-to-BLAST reranking, lookup/verified latency decomposition, DRAG graph construction, functional enrichment, and literature support analysis. The public repository tracks source code, benchmark task definitions, paper source, and compact result summaries; large local data files, model checkpoints, Chroma indexes, BLAST databases, and per-query JSON traces are regenerated from scripts rather than versioned. The reported vector lookup timings exclude model cold start, query embedding generation, LLM generation, and network calls. This choice isolates the indexed retrieval stage after embeddings are resident or precomputed.

BioRAG-DRAG is intended for research and evidence organization, not autonomous biomedical decision-making. The system can help retrieve, rank, and package evidence for a human or an auditable agent workflow, but it should not be used as a standalone clinical or experimental authority. The design deliberately keeps BLAST, curated annotations, graph evidence, and literature traces visible so that downstream users can inspect which route supported each claim.

## 10 Conclusion

BioRAG-DRAG provides a unified local retrieval layer for biomedical agents by combining neural sequence-text retrieval, classical biological verification, and evidence graph expansion. The current results show that vector retrieval is useful as a coarse multimodal candidate layer in a local sequence-window workflow, while BLAST remains essential for alignment-grounded verification. Hybrid DRAG graphs preserve broad vector connectivity and add typed biological evidence paths, making the system better suited to agent workflows than any single retrieval route alone. This positions BioRAG-DRAG as a practical engineering foundation for local-first biomedical agents, with biological graph analysis offering a promising but still exploratory path toward stronger biological interpretation.

## A Candidate-BLAST Ablation

Table 10 reports the full vector-to-candidate-BLAST budget sweep. Each query is vector-searched once at the maximum budget; smaller budgets use prefixes of that candidate list, so this isolates candidate-pool quality and candidate-BLAST verification rather than per-budget vector lookup latency. This table supports the engineering hypothesis that vector search can provide candidate pools for later alignment verification, but it is not treated as a main retrieval condition.

**Table 10:** Vector-to-candidate-BLAST budget sweep on the 100-query sequence-window stress test.

| Candidate budget | Exact Hit@10 | Exact MRR | Biological Hit@10 | Biological MRR | Candidate Bio Recall@N | Candidate BLAST latency |
| --- | --- | --- | --- | --- | --- | --- |
| 10 | 0.5000 | 0.3945 | 0.8283 | 0.8196 | 0.8283 | 172.0 ms |
| 25 | 0.5400 | 0.4517 | 0.8485 | 0.8394 | 0.8485 | 133.1 ms |
| 50 | 0.5700 | 0.4898 | 0.8687 | 0.8687 | 0.8788 | 131.6 ms |
| 100 | 0.6100 | 0.5151 | 0.8990 | 0.8990 | 0.9293 | 138.1 ms |
| 200 | 0.6500 | 0.5377 | 0.9293 | 0.9293 | 0.9596 | 146.2 ms |

**Table 11:** DRAG graph structure on 10k sequence-window graph views.

| Modality | Graph | Nodes | Edges | Components | Communities | Modularity | Key biological signal |
| --- | --- | --- | --- | --- | --- | --- | --- |
| DNA/cDNA | vector-only | 579 | 19,344 | 2 | 9 | 0.1741 | IGKV community: 50/66, enrichment $\times 6.178$ |
| DNA/cDNA | BLAST-only | 579 | 995 | 219 | 220 | 0.9383 | compact IGHV/IGKV/IGLV alignment modules |
| DNA/cDNA | hybrid | 579 | 19,592 | 2 | 7 | 0.1739 | IGKV 51/60, enrichment $\times 6.9317$; IG_J and IG_D modules |
| Protein | vector-only | 2,623 | 31,794 | 2 | 16 | 0.3179 | pgl 51/306 $\times 4.4158$; yfbR 42/226 $\times 11.6062$ |
| Protein | BLAST-only | 2,623 | 7,572 | 421 | 447 | 0.9721 | compact but fragmented alignment components |
| Protein | hybrid | 2,623 | 35,549 | 1 | 12 | 0.3779 | pgl 71/519 $\times 3.6245$; yfbR 42/63 $\times 41.6349$ |

**Table 12:** Vector-neighborhood enrichment against random neighbors.

| Target | Match | Vector Hit@10 | Random Hit@10 | Vector P@10 | Random P@10 |
| --- | --- | --- | --- | --- | --- |
| Protein windows | GN= / gene symbol | 0.4300 | 0.0200 | 0.1710 | 0.0020 |
| DNA/cDNA windows | gene ID | 0.6900 | 0.0600 | 0.2460 | 0.0060 |
| DNA/cDNA windows | gene symbol | 0.7188 | 0.0625 | 0.2563 | 0.0063 |
| DNA/cDNA windows | any identity | 0.7100 | 0.0600 | 0.2570 | 0.0060 |

**Table 13:** GO and Reactome enrichment of DRAG graph communities.

| Graph | GO annotated nodes | Reactome annotated nodes | Top GO signal | Top Reactome signal |
| --- | --- | --- | --- | --- |
| DNA/cDNA vector-only | 155 | 68 | proton transmembrane transport, 11/52, $q = 0.000113$ | Metabolism, 12/26, $q = 0.000015$ |
| DNA/cDNA BLAST-only | 155 | 68 | immunoglobulin mediated immune response, 11/11, $q = 0.000019$ | no mapped top term |
| DNA/cDNA hybrid | 155 | 68 | proton transmembrane transport, 11/50, $q = 0.000088$ | Metabolism, 12/26, $q = 0.000015$ |
| Protein vector-only | 63 | 213 | intracellular protein localization, 6/8, $q = 0.000516$ | metabolism of steroid hormones, 6/26, $q = 0.000452$ |
| Protein BLAST-only | 63 | 213 | postsynaptic membrane, 5/5, $q = 0.000042$ | GPCR downstream signalling, 9/25, $q = 0.000126$ |
| Protein hybrid | 63 | 213 | focal adhesion, 5/8, $q = 0.000539$ | TP53 regulates metabolic genes, 7/42, $q = 0.000223$ |

**Table 14:**
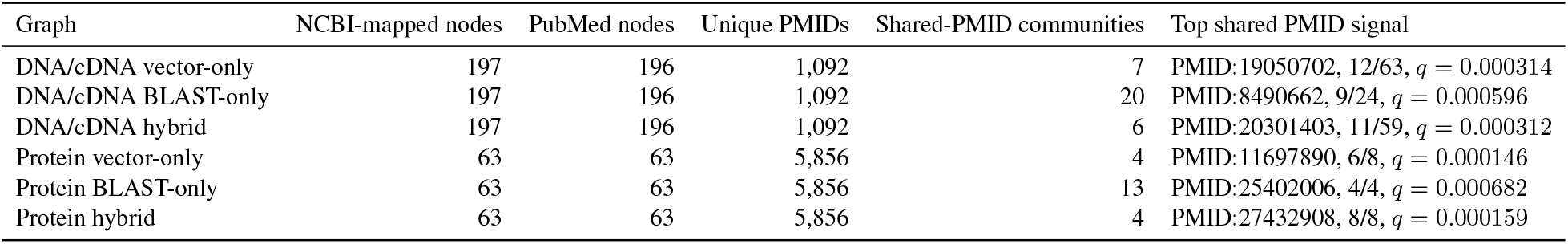
Shared PubMed evidence in DRAG graph communities.

| Graph | NCBI-mapped nodes | PubMed nodes | Unique PMIDs | Shared-PMID communities | Top shared PMID signal |
| --- | --- | --- | --- | --- | --- |
| DNA/cDNA vector-only | 197 | 196 | 1,092 | 7 | PMID:19050702, 12/63, $q = 0.000314$ |
| DNA/cDNA BLAST-only | 197 | 196 | 1,092 | 20 | PMID:8490662, 9/24, $q = 0.000596$ |
| DNA/cDNA hybrid | 197 | 196 | 1,092 | 6 | PMID:20301403, 11/59, $q = 0.000312$ |
| Protein vector-only | 63 | 63 | 5,856 | 4 | PMID:11697890, 6/8, $q = 0.000146$ |
| Protein BLAST-only | 63 | 63 | 5,856 | 13 | PMID:25402006, 4/4, $q = 0.000682$ |
| Protein hybrid | 63 | 63 | 5,856 | 4 | PMID:27432908, 8/8, $q = 0.000159$ |

## B Exploratory DRAG Biological-Structure Results

This appendix reports exploratory biological-structure analyses that are useful for hypothesis generation but are not used as main-text evidence for biological mechanism. The main text includes compact parent-collapse, k-mer/Jaccard, and degree-preserving null controls; the enrichment tables below should be read in light of those controls.

